# Strength-mass scaling law governs mass distribution inside honey bee swarms

**DOI:** 10.1101/2022.03.11.484032

**Authors:** Olga Shishkov, Claudia Chen, Claire Allison Madonna, Kaushik Jayaram, Orit Peleg

**Affiliations:** Biofrontiers Institute University of Colorado Boulder, Boulder, CO 80309, USA; Department of Computer Science University of Colorado Boulder, Boulder, CO 80309, USA; Santa Fe Institute, Santa Fe, NM 87501, USA

## Abstract

To survive during colony reproduction, bees create dense clusters of thousands of suspended individuals. How can this swarm, which is orders of magnitude larger than the size of an individual, maintain mechanical stability? We hypothesize that the internal structure in the bulk of the swarm, about which there is little prior information, plays a key role in mechanical stability and thermoregulation. Here, we provide the first-ever 3D reconstructions of the positions of the bees in the bulk of the swarm using x-ray computed tomography. We find that the mass of bees in a layer decreases with distance from the attachment surface. By quantifying the distribution of bees within swarms varying in size (made up of 4000-10000 bees), we find that the same power law governs the smallest and largest swarms, with the weight supported by each layer scaling with the mass of each layer to the ≈ 1.5 power. This arrangement ensures that each layer exerts the same fraction of its total strength, and on average a bee supports a lower weight than its maximum grip strength. This illustrates the extension of the scaling law relating weight to strength of single organisms to the weight distribution within a superorganism made up of thousands of individuals.

## 1 Introduction

When thousands of individual insects self-assemble into one coherent group, the resulting superorganism can perform functions that individuals cannot. Prominent examples include fire ant rafts and towers keeping a colony afloat during a flood and helping them get to dry land (1; 2), army ant bridges shortening a path of a marching colony, and their bivouacs forming a temporary home for their queen and brood (3; 4), and Western honey bees gathered in swarms to keep their queen safe while they search for a new hive (5). In these aggregations, the majority of the insects reside in the bulk of the structure, and could deeply affect its mechanics. Since it is not possible to see the individuals inside the opaque, dense aggregation, studies of these aggregations usually rely on observations of the individuals on the surface of the aggregations (5; 6). Some studies have used two-dimensional x-ray projections of the aggregations (2), but this method cannot quantify the distribution of individuals in all three dimensions. To address this gap, we use x-ray computed tomography (CT) to visualize the 3D arrangement of Western honey bees (*Apis mellifera*) in swarms, calculate how the load is distributed between bees, and use a scaling law to explain the reason for this weight distribution.

A honey bee swarm is a cluster made up of a queen bee and thousands of workers that hangs outside for hours to days while the workers scout for hives (7). Swarming is a precarious step in the life of a honey bee colony: if the queen or too many of the worker bees do not make it to their new hive, the entire colony will be lost (7). Bees attach to one another to support the weight of the entire swarm and keep the colony cohesive. Hence, the mechanical stability of these swarms plays an important role in the survival of the colony. So far, the arrangement of bees inside swarms and how a swarm’s weight is distributed between bees has been unknown. Previous studies on honey bee swarm structures investigate how the swarm shape changes in response to environmental perturbations. For instance, bees flatten the swarm to keep it stable if its attachment surface shakes (5), and respond to changing weather by packing the swarm more densely in the cold and spreading out in the heat (6; 8; 9).

To interpret the spatial distribution of bees measured by the x-ray CT, we hypothesize that a honey bee swarm is a superorganism that can be described with scaling laws, analogous to those that apply to individual organisms in nature. A number of scaling laws that relate the size of individuals to their physical properties have been established in biology (10; 11; 12). These laws are derived from dimensional analysis and can be verified experimentally. For instance, the metabolic rate of animals scales with body mass to the three-fourths (or two-thirds) power (10; 13), running speed scales with body mass to the one-sixth power (14; 15), and the strength of weightlifters scales with body mass to the two-thirds power (16). Just as an animal is an organism made up of cells, a swarm is a superorganism made up of bees, and similar scaling laws could apply to its physical properties.

In this study, we find a universal scaling law that applies to the arrangement of bees and weight distribution inside both small and large swarms. Regardless of swarm size, the weight supported by each horizontal layer of a swarm scales with the mass of the layer to the ≈1.5 power. This weight distribution assures that each swarm layer is using the same percentage of its maximum strength. We describe our methods for taking CT scans of swarms in Section 2. We show that the weight distribution inside the swarm obeys a power law in Section 3, and present a mathematical model explaining the benefits of the distribution of bees in the swarm scaling with this particular power law in Section 4. Finally, we discuss the results and conclude in Section 5.

## 2 Materials and methods

In this section, we provide a succinct decription of our methodology, including the honey bee swarms, our experimental setup, and our procedure for obtaining and and analyzing CT scans. See the Supplementary Methods for a more detailed description of the methodology.

### 2.1 Experimental setup

In our study, each swarm consists of a different number of worker bees (4000-9700 individuals) and a caged queen bee. We collected data from 11 swarms, providing 57 CT scans in total. We set up the swarms in a laboratory lit by ambient indoor lighting at an ambient temperature of 21°C. The swarm hangs from an attachement surface, a 33 cm diameter wood disk. The queen is attached to the center of the attachment surface, and the workers gather around her. We measure the weight of the swarms before taking CT scans, and calculate the number of bees in a swarm by dividing the mass of the swarm by the average weight of a bee.

We set up the swarm between the x-ray emitter and detector of a JPI DynaVue: Digital X-Ray and Fluoroscopy in One System, 17.7 ±0.7 cm from the detector. The emitter and detector are fixed 90 cm apart. The schematic in **Figure 1(a)** shows the swarm between the x-ray emitter and detector. We define the x-axis horizontal and parallel to the x-ray detector plane and the y-axis parallel to the x-ray emitter - detector line, so that the attachment surface and the xy-plane are coplanar. We define the z-axis in the direction of gravity, with the origin at the center of the attachment surface where the queen cage is attached. We center the origin of the axes in the field of view of the x-ray detector.

**Figure 1:**
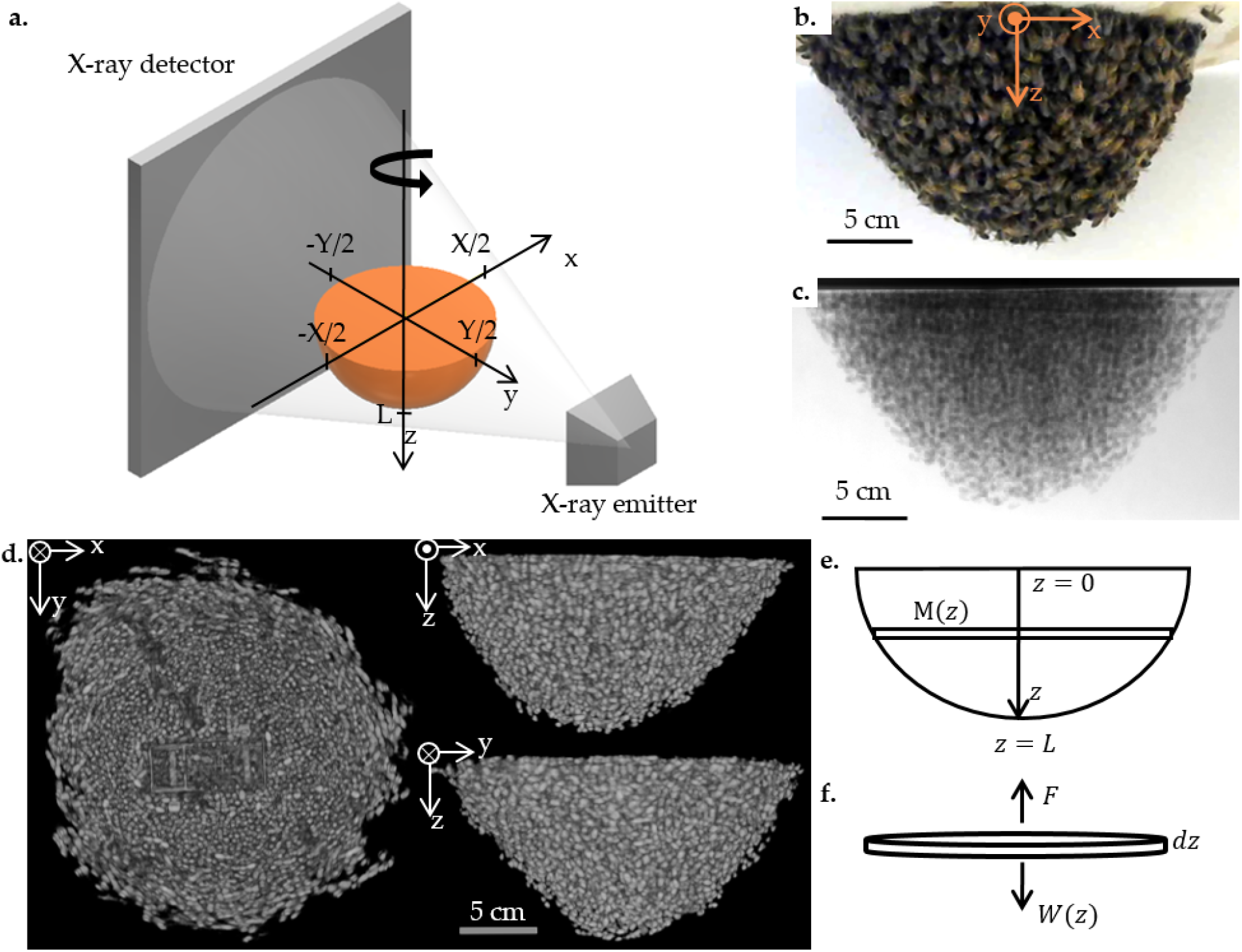
(a) Schematic of x-ray CT setup. The swarm hangs from a rotating board between the x-ray emitter and detector. As the swarm rotates about the z-axis, the emitter generates x-rays and the detector saves the resulting projections. (b) Photograph of a 753 g swarm taken right before a CT scan. (c) Sample projection of the CT scan of the swarm in (b). Darker gray pixels represent more bees in the path of the x-ray beam. (d) Volume rendering of a reconstruction of the swarm in (b), shown from three different orientations: top view, right below the attachment surface (left), front view (top right), and right-side view (bottom right). Brighter voxels represent a higher bee mass, and black voxels represent no bees. See Supplementary video V2 for a video of the rotated reconstruction. (e) Schematic of the swarm with a vertical coordinate *z*. The swarm is made up of layers with mass *M*(*z*) and thickness *dz*. (f) Free body diagram of a *dz* thick layer of the swarm. The force each layer of bees exerts to support the swarm, *F*, is equal to the weight of the layers underneath it, *W*(*z*).

We motorize the swarm with a stepper motor and use a MATLAB script to continuously rotate the stepper motor at 7.2°/sec, and save the motor position and x-ray parameters. The motor completes a full rotation for one CT scan in 25 seconds. While the motor rotates, we use customized proprietary software by JPI to acquire x-ray projections at 15 frames per second with x-ray parameters 90-95 kV and 20 mA. As a visual illustration, we show a photograph of the swarm hanging in front of the x-ray detector in **Figure 1(b)**, a sample projection of the same swarm in **Figure 1(c)**, and a sample set of projections in Supplementary video V1.

### 2.2 CT scan processing

To perform the 3D reconstructions, we first find the rotation angle of each projection, measure the length of the side of a voxel and the center coordinate, and subtract a blank background image to reduce noise. We then reconstruct each data set with the Matlab TIGRE toolbox, using the FDK algorithm and Hamming filter (17). See SI Section 11.4 for a more detailed description of the following processing and Supplementary Table T1 for a table of all the variables we use in our analysis. Each resulting reconstruction consists of a 3D matrix, *Ĩ*(*x*, *y*, *z*), of the brightness values in each voxel.

We set the bounds of the reconstructions to encapsulate the swarm and a small amount of empty space around it (unless the swarm reaches the edge of the field of view of the x-ray). The x-axis is bounded from −X/2 to X/2 and the y-axis is bound from -*Y*/2 to *Y*/2, as shown in **Figure 1(a)**. The z-axis ranges from 0 to the length of the swarm L, where 7.7 ≤ *L* ≤12.5 cm. X and Y are visually defined to encapsulate all non-zero CT brightness values (i.e., the physical shape of each swarm). We label all of the voxels that are encapuslated by the swarm and find its surface boundary as described in SI Section 11.4.

The raw reconstructions, *Ĩ*(*x*, *y*, *z*), have a cupping artifact throughout the swarm, likely from beam hardening (18), and noise that is caused by the short current and exposure time used to take the projections. Postproccesing to eliminate these artifacts (see SI Section 11.4 for details) results in a brightness matrix *I*(*x*, *y*, *z*) that is linearly correlated to the mass of bees in each voxel. We show three views of a sample reconstruction in **Figure 1(d)**, highlighting the radially symmetric shape of the swarm about the z-axis and the outlines of individual bees.

### 2.3 Converting brightness to mass and densities

To convert the brightness values *I*(*x*, *y*, *z*) into a measure of the mass of bees in each voxel, we use an external measure of the mass of the entire swarm, *M_swarm_* (see SI Section 11.3 for scale specs). Assuming the brightness, I, is linearly correlated with the density of bees, the following relation provides an estimated mass per voxel:

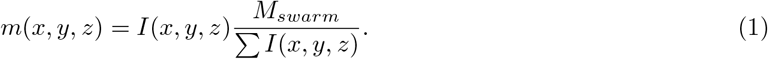

With the mass of each voxel and the boundary of the swarm at hand, we can define the density of bees *ρ*(*z*) in each layer (a one-voxel thick xy-slice). The length of the side of each voxel is *s*, and *N_B_* (*z*) is the number of voxels in a layer that are within the swarm boundary. Meausring *ρ*(*z*) shows how densely packed the bees are from the attachment surface to the swarm tip:

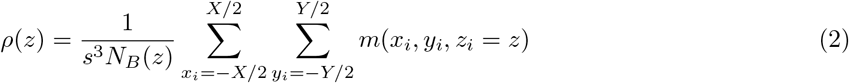

### 2.4 Mass and forces per layer

We use the matrix of the mass in each voxel, *m*(*x*, *y*, *z*), to analyze the weight distribution within the swarm by measuring the mass of each layer (a one-voxel thick xy-slice) and the weight supported by each layer.

The schematic in **Figure 1(e)** highlights a layer of the swarm. The mass of bees per unit distance in each layer, *M*(*z*), is the sum of the mass of each voxel in that layer divided by the thickness of the layer, *s*:

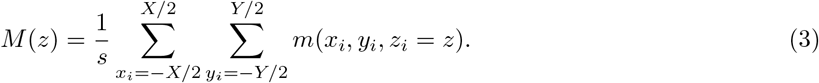

As in the free body diagram in **Figure 1(f)**, the force that each layer of the swarm exerts upwards is equivalent to the weight that the layer supports. The weight that each layer has to support, consisting of the weight of the bees in that layer and all of the layers underneath it, *W*(*z*), is:

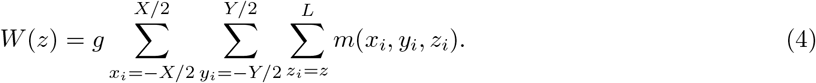

where *g* is the graviatational acceleration.

e discuss the implications of the trends in *M*(*z*) vs *W*(*z*) in Section 3. When applying curve fitting, we neglect data for *M*(*z*) and *W*(*z*) at the top and bottom edge: 1 cm at the top of the swarm, where *M*(*z*) has not yet reached its peak, and the few bees in the tip of the swarm, where *W*(*z*) ≤5 g. This allows us to limit our analysis of the weight distribution of the swarm to the trends in the bulk of the swarm and neglect outlying data from edge effects.

Finally, we measure the mean force each bee supports in a layer, *F_bee_*, assuming that each bee in a layer bears an equal fraction of the weight *W*(*z*). Since the voxel edge length is smaller than the length of a bee, we obtain the mass of individuals available to support the bees beneath them with a moving sum over the voxels that span the length of an individual bee, *l* = 1.5 cm long. We then divide *W*(*z*) by the mass of supporting bees:

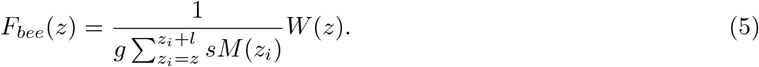

### 2.5 Maximum force per bee

We measure the maximum force that a bee can support by tying an elastic band to two bees, getting the bees to hold on to one another, and pulling them apart until they disconnect. We measure the initial length and final length of each elastic band and then use Hooke’s law to measure the force between the bees. See SI, Section 11.2 for details.

## 3 Results

In this section, we discuss 57 reconstructions of 11 swarms. We show three sample volume renderings of reconstructions of example swarms with small, median, and large masses in **Figure 2(a)**, and a reconstruction rotated to show the base, the tip, and different sides of the swarm in Supplementary video V2. For the rest of this paper we consider the bulk properties of the swarm, leaving the positions of individual bees a subject for future study.

**Figure 2:**
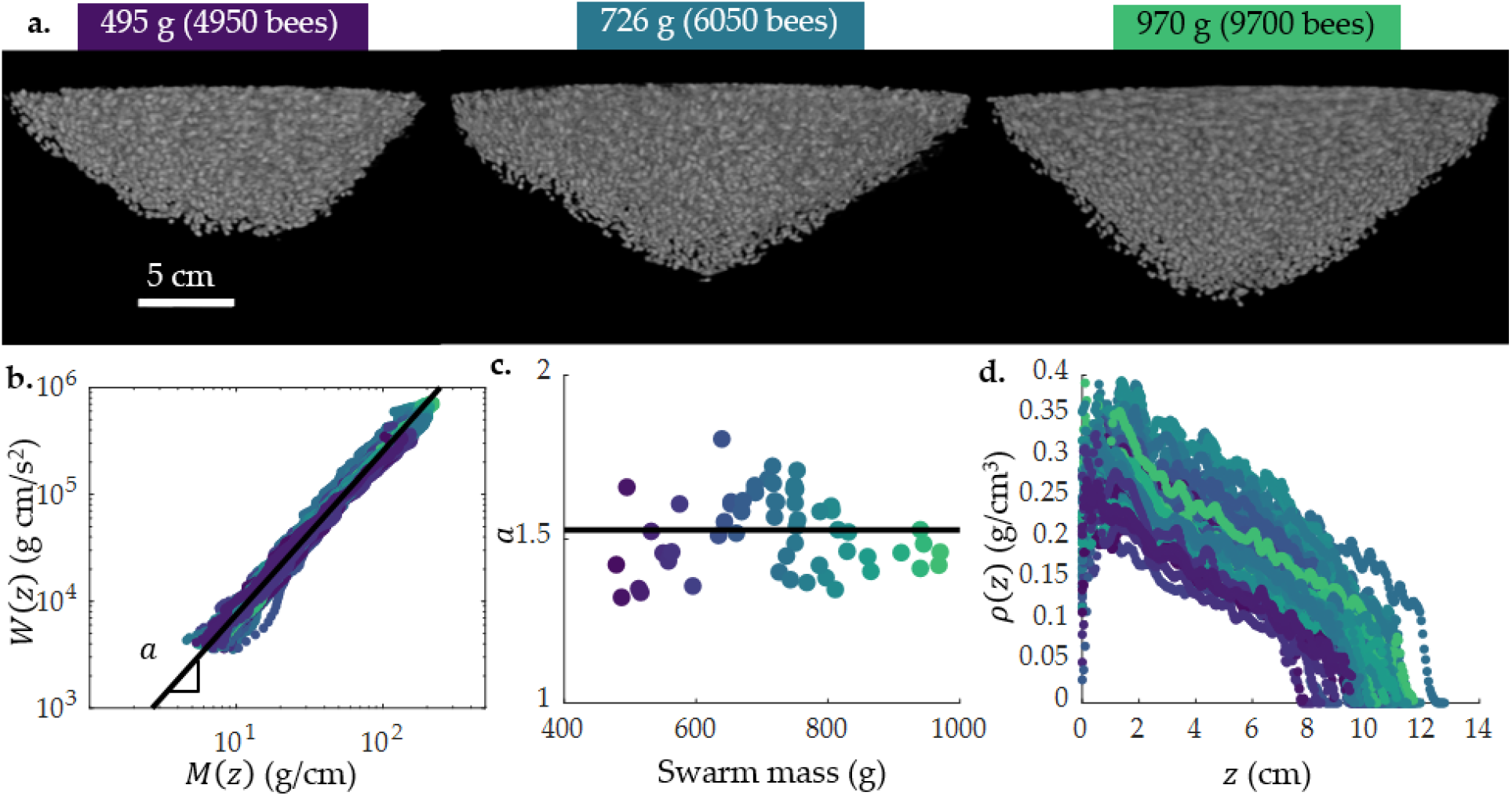
(a) Volume rendering of three different swarm reconstructions, in order of increasing mass: 495 g, 726 g, and 970 g. (b) Experimental data from all 57 reconstructions showing the relationship between weight underneath a layer, *W*(*z*), and mass of bees in that layer, *M*(*z*). Plotting *W*(*z*) and *M*(*z*) on a log-log scale shows that this relationship scales with the same power law for swarms weighing 480 to 970 g: *W*(*z*) = *CM*(*z*)^a^, *a* = 1.53 ± 0.12. (c) Plotting *a* vs. the swarm mass shows that *a* is independent of swarm mass. (d) The density of bees in a swarm, *ρ*(*z*), for all 57 reconstructions as a function of distance from the attachment surface to the tip of the swarm, *z*. In (b) - (d), each color repesents a different swarm, colored according to the mass: purple represents lighter swarms, and green represents heavier swarms.

### 3.1 Weight distribution inside swarms

How do bees distribute themselves throughout the swarm to support its weight? To answer this question, we calculate the mass of bees in each layer of the swarm, *M*(*z*), as in **Eq. 3** and the weight supported by those layers, *W*(*z*), as in **Eq. 4**. We plot *W*(*z*) vs. *M*(*z*) on a log-log scale in **Figure 2(b)** for swarms weighing 480 - 970 g (containing 4000 - 9700 bees). Despite the varying swarm sizes, the *W*(*z*) vs. *M*(*z*) curves all fall on a universal line, suggesting that swarms have the same fundamental structure at small and large scales. To further characterize this scaling law, we apply a linear best fit to each curve. The slope of this fit is the exponent *a* in the power law relating *M*(*z*) to *W*(*z*):

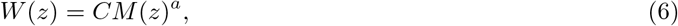

where *a* = 1.53 ± 0.12 and the constant *C* = 247 ± 110. We plot *a* for each swarm mass in **Figure 2(c)** and find that there is no trend in a as a function of the swarm mass, further supporting that weight distribution does not depend on the swarm mass.

**Figure 2(b)** shows that layers with the largest *M*(*z*) support the largest swarm weight *W*(*z*) (with *W*(*z*) = *gM_swarm_* at *z* = 0, where the top layer is supporting the entire swarm). Correspondingly, the three sample reconstructions in **Figure 2(a)** show that the area of swarm layers near the attachment surface is greater than the area of the layers near the tip of the swarm. To quantify the effect of packing density on layer mass, we plot the density of each layer of the swarm vs. z-coordinate for all 57 reconstructions using Eq. 2 in **Figure 2(d)**. The bees are most densely packed at the top of the swarm, and the density decreases towards the tip of the swarm, confirming that most of the bees are found near the base of the swarm.

To quantify the arrangement of swarm layers from the attachment surface to the tip of the swarm, resulting in the power law described in Eq. 6, we plot *M*(*z*) vs. *z* in **Figure 3(a)**. *M*(*z*) is high near the attachment surface, *z* ≤ 2 cm, and then decreases with *z* toward the tip of the swarm. This is visible in x-ray projections of the swarm, such as the one in **Figure 1(c)**. A darker shade of gray represents more bees between the x-ray emitter and x-ray detector, and white represents no bees. In this projection, there is a wide layer of darker gray representing densely packed bees near the attachment surface, followed by narrower layers of lighter gray representing less densely packed bees towards the swarm tip. The large mass of bees in the topmost layer creates a stable support structure for the bulk of the swarm, distributing the load between the bees.

**Figure 3:**
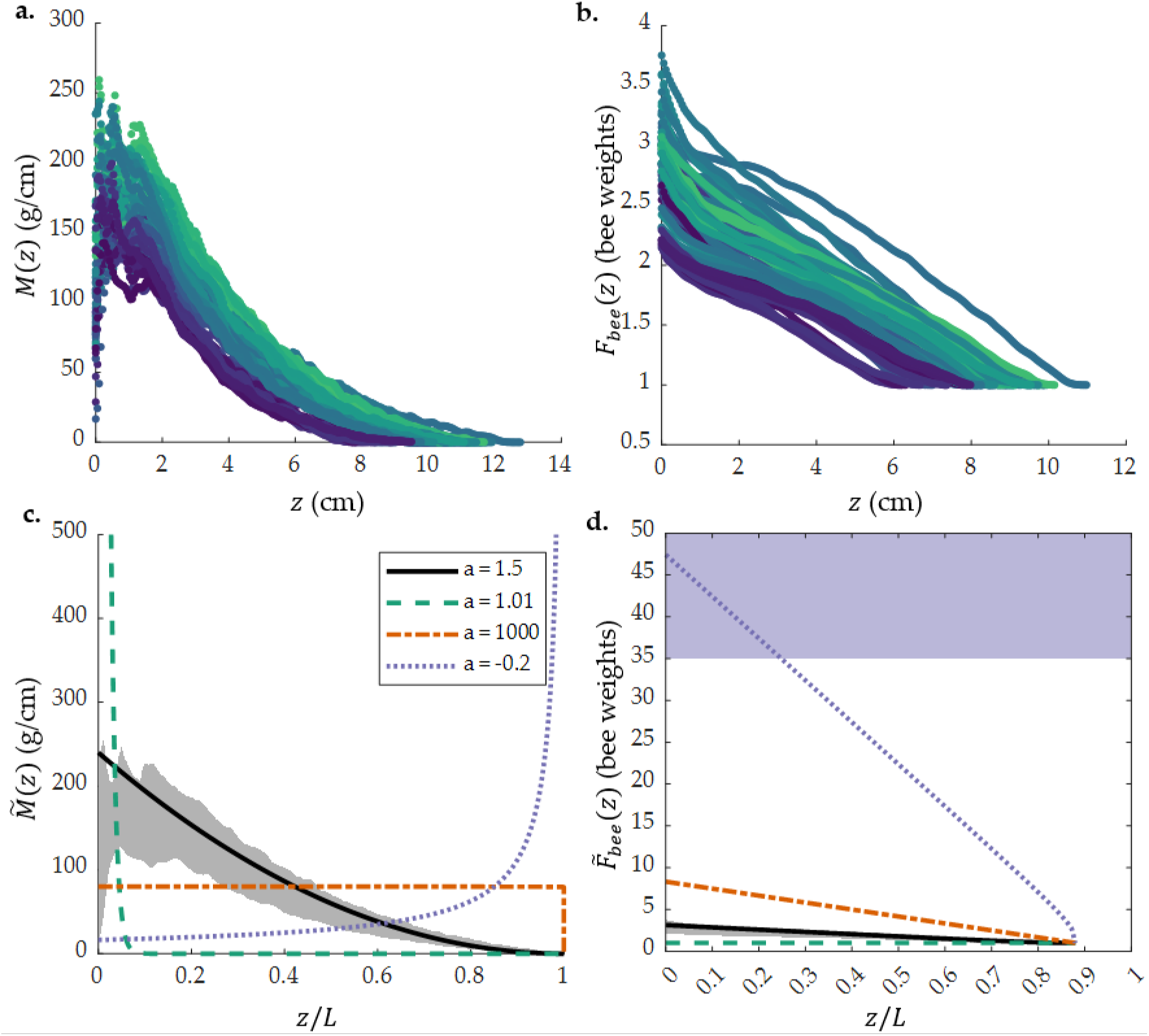
(a) Mass of swarm layers, *M*(*z*), plotted for all 57 reconstructions. *M*(*z*) peaks and then decreases with the *z* coordinate. (b) Mean force supported by a bee in a layer, *F_bee_*(*z*), vs *z*. The maximum force per bee is 3.8 bee weights, and the force decreases along the *z* axis. The lowest possible *F_bee_*(*z*) is one bee weight - a bee only supporting its own weight. (c) Theoretical mass distribution between the layers of a of a 1000-g, 12.5-cm long swarm with varying exponent a, with normalized z-coordinate *z*/*L* on the x-axis and mass per layer 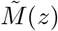 on the y-axis. The shaded gray region is the range of values for the experimental data in (a). (d) Theoretical mean force per bee for a 1000-g, 12.5-cm long swarm with varying exponent *a*, with *z*/*L* on the x-axis and force per bee 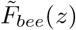 on the y-axis. The shaded gray region is the range of values for the experimental data in (b); the shaded purple region for 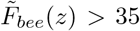 bee weights represents the maximum number of bees that a single bee can support. In (c) and (d), the solid black line corresponds to *a* = 1.5: the experimentally measured mass distribution, with more massive layers near the attachment surface (*z*/*L* = 0). Dashed green line corresponds to *a* = 1.01: a swarm with most of its mass concentrated in the top layer of the swarm, and very few bees found beneath that layer. Dash-dot orange line corresponds to *a* = 1000: a swarm in which all of the layers have equal mass. Dotted purple line corresponds to *a* = −0.2: a swarm in which the layers near the attachment surface have a lower mass than the layers at the bottom of the swarm.

How does this arrangement affect the forces experienced by individual bees in the swarm? If the weight supported by an individual bee is close to its limit, that bee will be more likely to break its bond when the swarm is mechanically perturbed. To investigate how much weight individual bees support, we calculate the mean force supported by a bee in a layer, *F_bee_*(*z*), as in Eq. 5. We plot *F_bee_* vs. *z* in **Figure 3(b)** and find that each bee supports a maximum of 3.8 bee weights. The weight supported per bee decreases from attachment board to swarm tip. Our experiments with pairs of bees show that one bee can support a maximum weight of 35 ±14 bees (see SI section 11.2). Hence, the bees bearing the weight of the swarm are not at their strength limit.

## 4 Scaling law of strength and layer mass

Our experimental data reveals a scaling law between the mass of a layer along the vertical coordinate, *M*(*z*), and the weight that it supports, *W*(*z*), namely: *W*(*z*) ~ *M*(*z*)^a^ with *a* ≈ 1.5. To better understand the physical mechanism that yields this scaling law, we derive the force balance equation of a layer of the swarm and solve for *W*(*z*). We then equate the analytical expression for *W*(*z*) with the experimentally determined scaling law, *W*(*z*) ~ *M*(*z*)^a^, to connect the swarm mass distribution to the exponent a and formulate the expressions for *M*(*z*) and *W*(*z*) in terms of a. We then consider a dimensional analysis of the strength of each layer of the swarm, *S*, or the maximum weight that it can support before the grip of the bees on one another breaks. As will be described in detail below, we find that *S* ~ M^1.5^, which is close to the experimentally determined *a* = 1.53. Deviation from this value increases the fraction of maximum strength exerted by different parts of the swarm.

### 4.1 Force balance model of the weight distribution in the swarm

We assume that the swarm is at quasi-equilibrium (the shape does not change although individual bees may move), that all of the bees in each layer contribute equally to supporting the weight of the bees underneath that layer, that the layer thickness is very small, and that the swarm is radially symmetrical about the z-axis. We use a cylindrical coordinate system with a vertical coordinate *z*, as shown in **Figure 1(e)**, and we consider layers of the swarm along the *z*-axis of thickness *dz*. Variables labeled with a tilde, as in 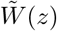, represent analytically derived expressions; variables without a tilde, as in *W*(*z*), represent values determined with power law fits to experimental data.

We begin our analysis by applying the force balance principle to each layer of a swarm. As shown by the free body diagram in **Figure 1(f)**, the force with which each layer of bees has to grasp the layer above it is equal to the weight of that layer and all of the layers underneath it: 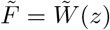. We express 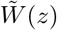 using the force balance equation (a continuous version of the discrete definition in Eq. 4.):

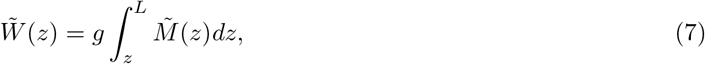

where the mass of bees per layer is 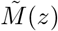, the swarm length is *L*, and *g* is the gravitational constant. Inspired by our experimental observation that the mass of the layers near the base is highest and the mass of the layers at the tip of the swarm is lowest in **Figure 3(a)**, we model 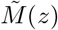 as a monotonically decreasing function of *z*. To keep the units consistent, we normalize the *z* coordiante by the length of the swarm:

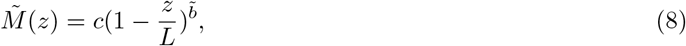

where the *c* factor in this expression ensures that the units of the mass per layer are mass/length, and 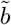 is an unknown exponent. Choosing this function form allows us to easily integrate the expression for 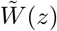 when we substitute 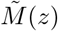 into it, set this force balance derivation for 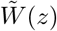 equal to the experimentally determined expression *W*(*z*) = *CM*(*z*)*^a^*, and compare the exponents *a* and 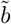. This expression for the layer mass of a swarm takes into account both changes in layer area, *Ã*(*z*), and layer density, 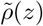, and could be expressed as 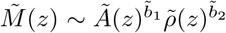 to separate the effects of the area and density. Both the density, as seen in **Figure 2(d)**, and the area of each layer, as seen in the reconstructions in **Figure 1(d)** and **Figure 2(a)**, decrease with the z-axis. In this analysis, we consider the area and density effects on the mass distribution with one exponent, 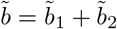.

To solve the expression for 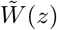, we substitute the expression for 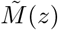, Eq. 8, into Eq. 7 and integrate. We then express 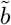 in terms of the experimentally determined *a* by equating this expression for 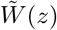 to the scaling law we observe in our experiments, Eq. 6, 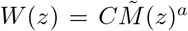. The exponent in the expression for 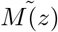, Eq. 8, is

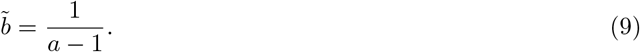

The weight supported by each layer is then:

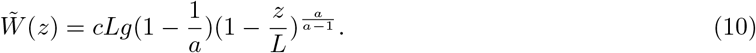

Next, we test how well our force balance model predicts the data by comparing the predicted value of 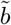 using the force balance to the value of *b* calculated using experimental fits. We first calculate 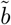 using the expression derived from the force balance, Eq. 9, and our experimental result for *a*, which yields 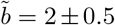. Second, we calculate *b* by applying a power law fit to the data in **Figure 3(a)**, *M*(*z*) ~ (1 – *z*/*L*)^b^, which yields *b* = 1.7 ± 0.2. See Supplementary Figure S5(a) for the *M*(*z*) vs 1 – *z*/*L* log-log plot. We calculate the deviation of 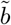 from *b*, 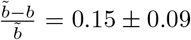, and plot the deviation of *b* from 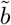 in Supplementary Figure S5(b) as a comparison for the individual CT scans. The values of *b* and 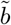 being on the same order of magnitude validates the model and allows us to compare 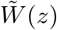 to a maximum strength of each layer, which we find with dimensional analysis in the following section.

### 4.2 Strength of a swarm layer and individual bees

The strength of the layer, 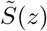, or the maximum weight that it could support, can be greater than or equal to 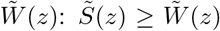. If the weight of the bees underneath a layer were to exceed its strength 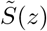, the layer would not be able to support the weight of those bees, and the swarm would break apart. We perform a dimensional analysis on the strength of each layer to find the relationship between the mass of a layer and its maximum strength, 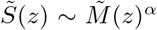. Force is proportional to mass, which is proprtional to volume, or a length cubed, so a layer’s strength scales with length cubed, 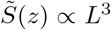. The mass of each layer, with units of mass/length, is proportional to an area, or a length squared, so 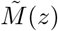 scales with length squared, 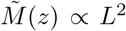. Thus, *α* must be 1.5 for 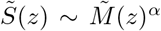 to be dimensionally correct. This is similar to the relaitonship between weightifting capacity and body weight in Ref. (16).

Estimating 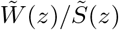 gives a measure of how much of its maximum strength each layer uses to hold up the rest of the swarm:

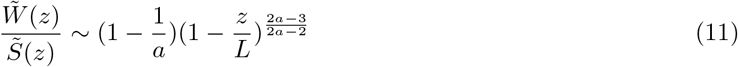

The average number of bees that a bee in a swarm layer supports, 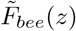, is equal to the mass of bees supported by a layer divided by the sum of the mass of bees in a layer of bees that has the thickness of the length of a bee, *l* ≈ 1.5, as a continuous version of the discrete equation in Eq. 5:

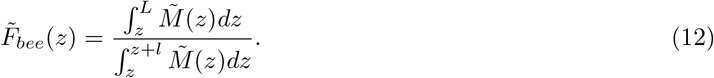

After integrating, we get an expression for 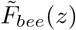:

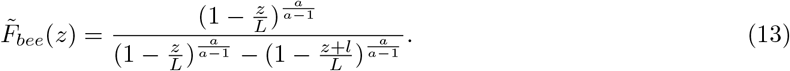

We then use the expression for 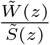, Eq. 11, and 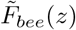, Eq 13, in the next section to evaluate how the force distribution in the swarm would change for swarms with different values of *a*.

### 4.3 Effect of *a* on the mass of each layer, the fraction of its maximum stregnth it uses, and the average force per bee

We now consider the effect of varying *a* on the mass and force distribution inside the swarm. To visualize the effect of *a* on the distribution of bees, we plot the mass per layer of a 1000-g, 12.5 cm long swarm, 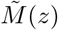 vs. *z*/*L*, with *a* = 1.5,1.01, 1000, and −0.2 in **Figure 3(c)** and the corresponding average force per bee, *F_bee_*(*z*) vs. *z*/*L* in **Figure 3(d)**. These values of *a* are example values for the four possible cases of mass distribution in the swarm. We then evaluate how these values of *a* affect the fraction of maximum strength each layer uses to support the layers underneath it using Eq. 11.

If *a* ≈ *α*, as we found in our experiments, layers with higher mass near the attachment surface support the less massive layers under them, as in the solid black line in **Figure 3(c)**. Correspondingly, **Figure 3(d)** shows 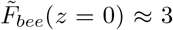 at the top of the swarm, and decreases towards the tip. The strength of each layer and the weight it supports are proportional to one another, 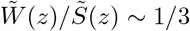, meaning that the fraction of maximum strength used by a layer is the same for all *z*. If 1 < a < α, the swarm approaches one massive layer of bees, as in the dashed purple line in **Figure 3(c)**. The dimensional analysis results in a very small fraction of the total strength used by this layer, 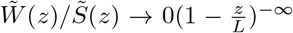. The force supported by each bee in **Figure 3(d)** shows 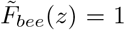 for the entire swarm, meaning that each bee only supports its own weight. This configuration would either require packing a large number of bees into one very dense or one very wide layer. A swarm with one very dense layer at the top would compress all of the bees; a swarm with one very wide layer would require a large surface area, which would put the swarm in danger from predators and changes in weather. Thus, despite a potentially lower fraction of strength used by the largest layer of bees, this configuration would put the swarm in danger by requiring a large surface area.

For values of *a* > α, as *a* → ∞, all the layers of the swarm have the same mass, as in the dash-dot red line in **Figure 3(c)**. The force per bee in **Figure 3(d)** shows 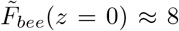 at the top of the swarm, 2.5 times that of the a =α configuration. In this configuration, the top layers use a higher percentage of their available strength than the lower layers, 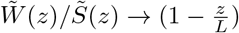. Thus, for large swarms, the bees that support the swarm would be under more strain, and the swarm would be more likely to break under external perturbation.

Finally, a < 0 (0 ≤ *a* ≤ 1 results in negative values for 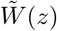) would suggest that the top layers of the swarm have a lower mass than the bottom layers, as in the dotted orange line in **Figure 3(c)**. This is not a realistic range of values for *a*, but we include it here as a demonstration of a potential mass distribution with the largest layers being on the bottom of the swarm. This configuration would put even more strain on the layers of bees at the top of the swarm, as smaller layers near the attachment surface have a smaller maximum strength. As *a* →0 on the *a* < 0 side, 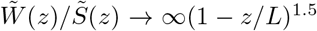, and bees in the top layers use a much greater fraction of their strength than bees in the bottom layers. Accordingly, the mean force per bee in **Figure 3(d)** exceeds the maximum bee grip strength of 35 bee weights, and the swarm could not support itself in this configuration.

The swarm configuration with *a* ≈ 1.5 uses the full strength of each layer and puts a lower strain on the bees than most other values of *a*, and avoids weight distributions that could expose a large number of bees to external danger.

## 5 Discussion

In this study, we obtained x-ray CT scans of western honey bee swarms and analyzed trends in the bulk distribution of bees. We find that, regardless of swarm size, bees are arranged such that the weight supported by each layer scales as the mass of the layer to the ≈ 1.5 power. This supports our hypothesis that scaling laws can be used to describe a superorganism such as a honey bee swarm, in a similar way to past descriptions of individual animals and plants in the past (10). Examples of scaling laws which have previously been used to understand collective behavior in groups of animals include studies relating the number of ants pulling an object to the speed of that object (19; 20), spatial and temporal correlation of motion in insect swarms (21), and size of fish groups over time (22; 23). Our scaling law relating the mass of layers to the weight they support is unusual in that, rather than relating the strength of a swarm to the number of individuals within the swarm, it relates the force distribution within the swarm to the arrangement of individuals inside it. Regardless of the number of individuals in the swarm, the same scaling law can describe its structure.

We found that bees are arranged to distribute the weight between each other without overloading individual bees, which can leave some bees free to scout for new hive locations. We primarily consider the bulk properties of the swarm, but x-ray CT data provides a 3D image of all of the bees inside the swarm, and there is a wealth of untapped data in our scans regarding how the bees are distributed within the layers. Our preliminary analysis of the internal structure of the swarm suggests that bees form a “scaffold” inside the swarm that leaves other bees free to move within the swarm and scout for new hives, as shown in the slices in Supplementary Figure S4 and Supplementary videos V3 and V4. Our future studies will investigate how bees might use this internal scaffolding to navigate throughout the swarm.

We focused this study on swarms at an ambient room temperature. Different ambient temperatures might result in swarms deviating from *a* ≈ 1.5 to adjust the swarm configuration for thermoregulation. Heinrich proposed an internal structure for swarms in cold or hot ambient environments in (9): homogenously packed bees within a dense shell in cold weather to generate heat, and air channels within the swarm for ventilation at hot temperatures. However, measurements of the shape change of the swarm in response to heating and cooling show that the swarm’s shape does not always directly correlate with temperature (8). X-ray CT provides an opportunity to see inside swarms in varying environmental conditions and investigate how the arrangement of bees changes to maintain a comfortable core temperature. Shape change in response to environmental conditions may also result in a tradeoff between stability and thermoregulation, with a different value for a.

The insight gained from this study on how honey bee swarms support their weight is both a fascinating addition to our understanding of the process by which honey bee colonies reproduce, and can also serve as a starting point for bioinspired engineering applications (24). A swarm can be thought of as a living material that can adjust to environmental conditions and repair itself when damanged. Engineering materials that similarly respond to their environment and self-heal might have applications in medicine, building and aircraft design, and smart sensors (25; 26). Biological swarms can provide design inspiration for novel robotic swarms (27). For instance, collectives of insect scale robots (28) hanging (29) from a building as they repair part of it may be inspired by the arrangement of bees in the swarm, ensuring that their assembly is stable. These practical applications in combination with quantitative ethological studies of the collective intelligence of these social insects could inform future studies of honey bee swarms.

## 6 Author contributions

O.S. and O.P. concieved the research study and designed the experiments; O.S., C.C., and C.M. performed the experiments; O.S. performed data analysis with the help of C.C. and C.M.; O.S., K.J., and O.P. designed the mathematical model; O.S., K.J., and O.P. wrote the paper; O.P. supervised the project.

## 7 Acknowledgements

This work was supported by the National Science Foundation (NSF) Physics of Living Systems Grant No. 2014212 (O.P.). Any opinions, findings, and conclusions or recommendations expressed in this material are those of the authors(s) and do not necessarily reflect the views of the NSF. We also acknowledge funding from the University of Colorado Boulder, BioFrontiers Institute (internal funds), the Interdisciplinary Research Theme on Multi Functional Materials and Autonomous Systems (O.P.). We would like to thank Profs. Dan Goldman, David Hu and L. Mahadevan for helpful discussions, Will Little, Andriy Nevgasymyi, and Gary Nave for assistance with the x-ray setup and software, Adrian Gestos for advice on CT reconstructions, Alexander Lawson for assistance with the experimental setup, Golnar Gharooni Fard for reading and commenting on the manuscript, Seneca Kristjondottir and Christopher Borke for beekeeping assistance, and Elizabeth Bradley, Karen McClune, and Sean Burns for hosting beehives.

