## Supplementary Info for "Strength-mass scaling law governs mass distribution inside honey bee swarms"

### 8 Supplementary tables

.

| Swarm trial | Date | Time | Swarm mass (g) |
|---|---|---|---|
| S03 | 10-May | 14:01 | 751 |
| S03 | 11-May | 13:15 | 719 |
| S03 | 11-May | 14:21 | 719 |
| S03 | 11-May | 14:42 | 716 |
| S03 | 11-May | 15:08 | 709 |
| S03 | 12-May | 15:29 | 690 |
| S03 | 12-May | 15:57 | 689 |
| S03 | 12-May | 16:08 | 689 |
| S04 | 24-May | 13:11 | 831 |
| S04 | 24-May | 13:22 | 815 |
| S04 | 24-May | 13:31 | 805 |
| S04 | 24-May | 13:39 | 806 |
| S04 | 24-May | 15:04 | 788 |
| S04 | 25-May | 12:15 | 753 |
| S04 | 25-May | 15:15 | 749 |
| S04 | 25-May | 16:06 | 748 |
| S04 | 26-May | 12:04 | 752 |
| S04 | 26-May | 13:14 | 754 |
| S04 | 26-May | 14:28 | 750 |
| S05 | 1-Jun | 14:00 | 722 |
| S05 | 1-Jun | 14:15 | 717 |
| S05 | 2-Jun | 11:30 | 653 |
| S05 | 2-Jun | 12:51 | 653 |
| S05 | 2-Jun | 16:34 | 639 |
| S09 | 21-Jul | 10:06 | 532 |
| S09 | 22-Jul | 10:11 | 550 |
| S10 | 29-Jul | 09:27 | 595 |
| S11 | 3-Aug | 10:50 | 643 |
| S11 | 4-Aug | 10:20 | 575 |
| S11 | 5-Aug | 10:30 | 495 |
| S12 | 10-Aug | 11:51 | 829 |
| S12 | 10-Aug | 12:56 | 750 |
| S12 | 11-Aug | 11:19 | 671 |
| S12 | 11-Aug | 12:05 | 669 |
| S12 | 12-Aug | 10:25 | 661 |
| S12 | 12-Aug | 11:38 | 634 |
| S13 | 23-Aug | 12:09 | 970 |
| S13 | 23-Aug | 12:55 | 968 |
| S13 | 24-Aug | 10:16 | 945 |
| S13 | 24-Aug | 12:17 | 940 |
| S13 | 25-Aug | 08:55 | 940 |
| S13 | 25-Aug | 10:23 | 911 |
| S14 | 25-Aug | 14:49 | 797 |
| S14 | 25-Aug | 15:47 | 787 |
| S14 | 26-Aug | 09:43 | 726 |
| S14 | 27-Aug | 10:39 | 487 |
| S14 | 27-Aug | 11:05 | 479 |
| S15 | 30-Aug | 13:29 | 865 |
| S15 | 30-Aug | 14:10 | 860 |
| S15 | 31-Aug | 10:11 | 811 |
| S15 | 31-Aug | 10:42 | 768 |
| S15 | 1-Sep | 09:42 | 743 |
| S15 | 1-Sep | 10:28 | 737 |
| S16 | 1-Sep | 14:56 | 558 |
| S16 | 1-Sep | 15:24 | 563 |
| S16 | 2-Sep | 10:41 | 513 |
| S16 | 2-Sep | 11:08 | 516 |

Supplementary Table T1: Summary of the dates, times and weights of swarms in CT scans used in this study.

| Symbol | Description | Units |
|---|---|---|
| $x$ | Coordinate parallel to x-ray detector plane | cm |
| $y$ | Coordinate along the detector – emitter axis | cm |
| $z$ | Vertical coordinate pointing down from center of swarm base | cm |
| $X/2$ | Maximum dimension of the x-coordinate | cm |
| $Y/2$ | Maximum dimension of the y-coordinate | cm |
| $L$ | Length of swarm | cm |
| $l$ | Length of a bee | cm |
| $M_{swarm}$ | Mass of the swarm | g |
| $g$ | Gravitational acceleration | cm/s$^2$ |
| $s$ | Length of the side of one voxel | cm |
| $N_B$ | Number of elements within the boundary of the swarm. | |
| $\tilde{I}(x,y,z)$ | Brightness values in each voxel of a reconstruction before postprocessing | |
| $I(x,y,z)$ | Brightness values after postprocessing | |
| $m(x,y,z)$ | Mass in each voxel | g |
| $\rho(z)$ | Average density of a layer of the swarm at vertical coordinate $z$. | g/mL |
| $F$ | Force exerted by a layer of bees to support the swarm | g cm/s$^2$ |
| $M(z)$ | Mass of bees in a layer of the swarm in a reconstruction | g/cm |
| $\widetilde{M}(z)$ | Mass of a layer of the swarm in the force balance model | g/cm |
| $W(z)$ | Total weight a layer supports in a reconstruction | g cm/s$^2$ |
| $\widetilde{W}(z)$ | Total weight supported by a layer of the swarm in the force balance model | g cm/s$^2$ |
| $\tilde{S}(z)$ | Maximum strength of a layer of a swarm | g cm/s$^2$ |
| $a$ | Exponent in the power law , $W(z) = C\ M(z)^a$, calculated by applying a power law fit to the $W(z)$ vs. $M(z)$ data | |
| $\alpha$ | Exponent in $\tilde{S}(z) \propto \widetilde{M}^{\alpha}$, found using dimensional analysis | |
| $C$ | Coefficient in the experimentally determined power law , $W(z) = C\ M(z)^a$ | g$^{0.5}$s$^{-2}$cm$^{2.5}$ |
| $\tilde{C}$ | Coefficient in the experimentally determined power law, estimated from the force balance model | g$^{0.5}$s$^{-2}$cm$^{2.5}$ |
| $\tilde{b}$ | Exponent in $\widetilde{M}(z) = \bar{\rho}L^2\left(1-\frac{z}{L}\right)^{\tilde{b}}$ , calculated from the experimentally determined $a$ | |
| $b$ | Same exponent as $\tilde{b}$, but calculated by applying a power law fit to the $M(z)$ vs. $1 - z/L$ data | |
| $F_{bee}(z)$ | Average number of bee weights supported by a bee in a layer measured from the experimental data | Bees |
| $\tilde{F}_{bee}(z)$ | Average number of bee weights supported by a bee in a layer estimated analytically | Bees |
| $\tilde{A}(z)$ | Area of a layer of the swarm in the force balance model, characterized by exponent $\tilde{b}_1$ | cm$^2$ |
| $\tilde{\rho}(z)$ | Density of a layer of the swarm in the force balance model, characterized by exponent $\tilde{b}_2$ | g/mL |
| $\bar{\rho}$ | Mean density of the entire swarm | g/mL |

Supplementary Table T2: Summary of the variables we use in this paper.

### 9 Supplementary figures

.

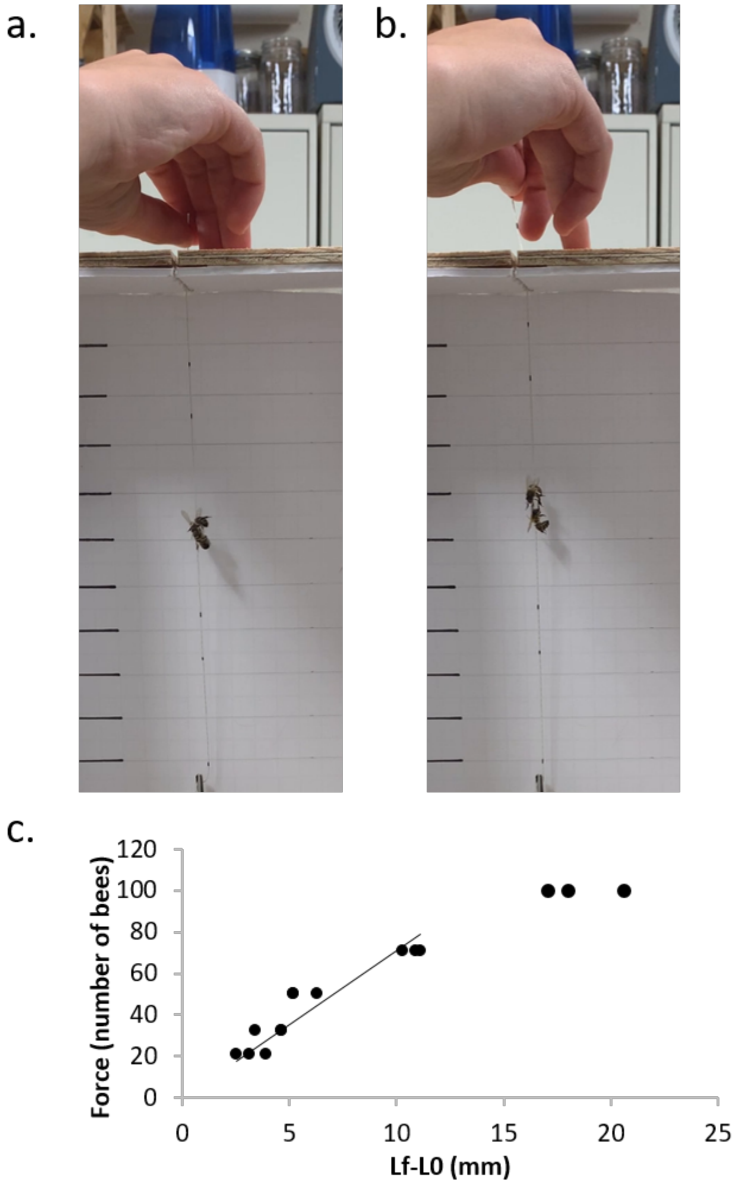


Supplementary Figure S1: (a) Two bees gripping one another before the elastic string is stretched. The elastic attached to the bottom bee is clamped in place, and we hold the elastic attached to the top bee. For each bee, we measure the initial string length $L0$. b) Two bees gripping each other while being stretched, right before they lose their grip. We measure the final length $Lf$ for each bee, and use Hooke's law to find the force on each bee. (c) Calibration data for the elastic string used to measure maximum bee grip strength.

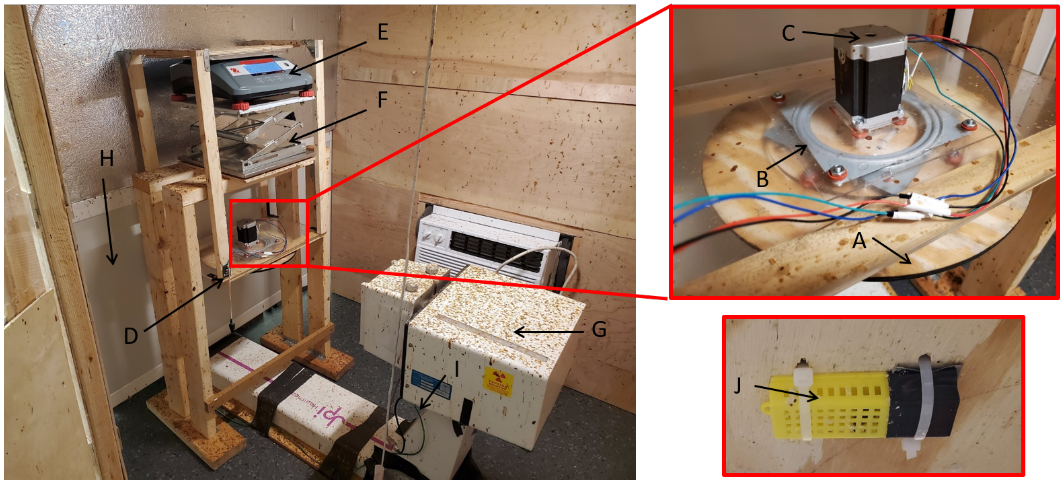


Supplementary Figure S2: Photograph of the experimental setup. (A) Attachment surface. (B) Turntable. (C) Stepper motor. (D) Motor controller. (E) Scale. (F) Positioning jack. (G) X-ray emitter. (H) X-ray detector. (I) Webcam (underneath x-ray generator). (J) Queen cage.

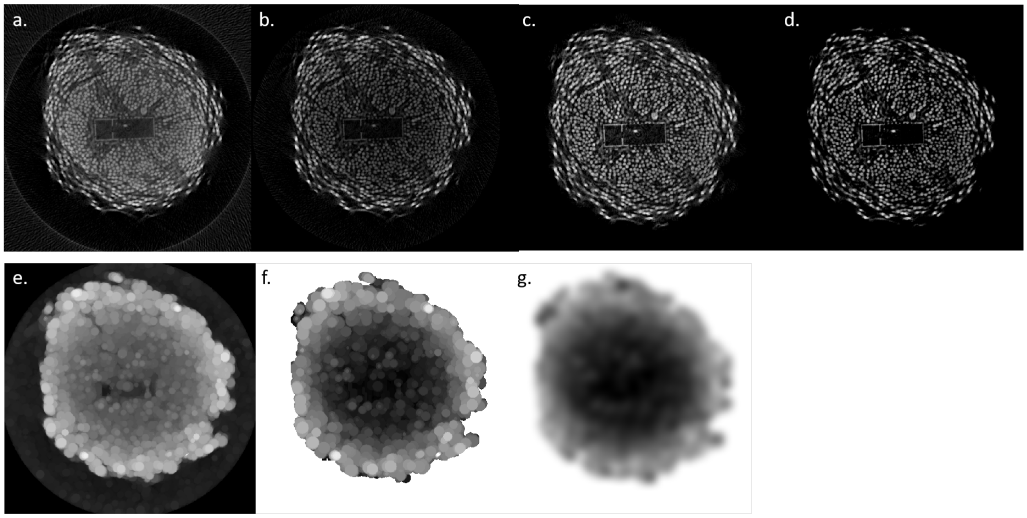


Supplementary Figure S3: (a) Raw slice before filtering and normalization. (b) Slice after removing noise outside swarm radius and performing top-hat filter for noise reduction. (c) Slice after normalization by normalization matrix in (g). (d) Final slice after background noise subtraction. (e) Dilated slice. (f) Dilated slice with the values outside the swarm boundary maximized to remove edge effects. (g) Normalization matrix created by blurring the slice in (f) with a Gaussian filter.

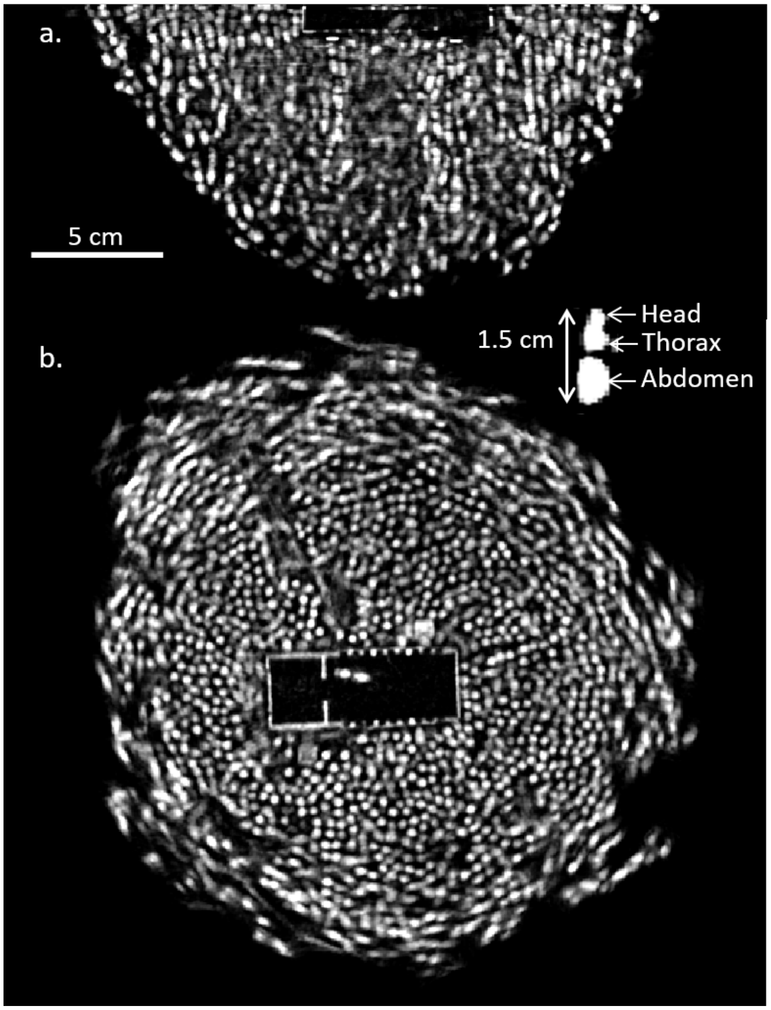


Supplementary Figure S4: (a) An 0.58-cm thick xz-slice taken at the centerline of the 753 g swarm in **Figure 1(b-c)**. Bees show up in this slices in two parts - a small circle for the thorax and a longer oval for the abdomen. Bright white represents bees that were stationary for the duration of the scan. Gray regions represent bees that could not be resolved due to the motion blur during the 25-second duration of the CT scan. Inset: a close-up slice of a bee with the head, thorax, and abdomen labeled. Scale bar is 5 cm, and applies to both (a) and (b). (b) An 0.58 cm thick xy-slice of the swarm, taken 0.29 cm from the attachment surface. The rectangular space in the center is the outline of the queen cage; the bee within that box is the queen. Round outlines are cross-sections of bees hanging vertically from attachment board.

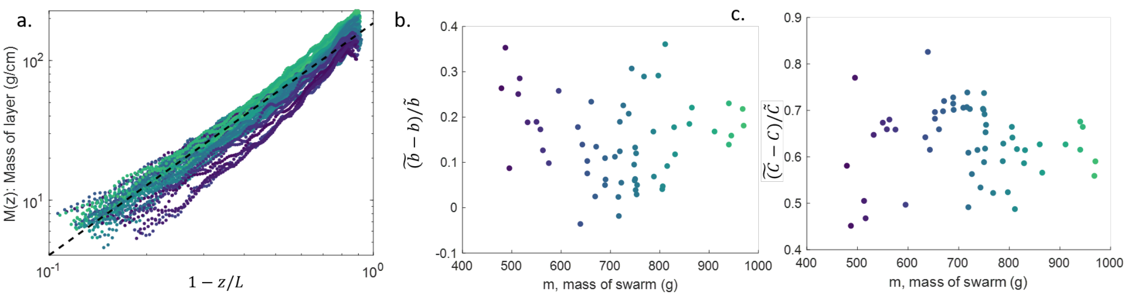


Supplementary Figure S5: (a) $M(z)$ vs. $1 - z/L$ plotted on a log-log scale. (b) Deviation of $b$ (calculated with a best-fit line to data in (a)) from $\tilde{b}$ (calculated from $b = 1/(a-1)$). (c) Deviation of $C$ (calculated with a best-fit line to data **Figure 2(c)**) from $\tilde{C}$ (calculated from the swarm parameters).

### 10 Supplementary videos

V1 All 374 raw projections of one full revolution of the swarm in **Figure 1(b-d)** for one CT reconstruction. The video is saved at 15 FPS for a total of 25 seconds per revolution.

V2 A 3D view of the outside of the swarm in **Figure 1(b-d)**, rotated to show the arrangement of bees at the attachment surface, the tip of the swarm, and the different angles of the swarm.

V3 Time lapse of the xz-slice of the swarm in Supplementary Figure S4 (a) with 25 s between frames. The bright regions, chains of bees, remain static during the scan while other bees move around them.

V4 Time lapse of the xz-slice of the swarm Supplementary Figure S4 (b) with 25 s between frames. The bright regions, chains of bees, remain static during the scan while other bees move around them.

### 11 Supplementary Info

#### 11.1 Swarm care

In our study, a swarm consists of 480 - 970 g of worker bees and a queen bee. We describe CT scan data from 11 swarms performed during May to August 2021.

We present data from a total of 57 scans gathered from 11 swarms. We show a summary of the scans collected for each swarm in Supplementary Table T1. We set up the swarms in a laboratory lit by ceiling lights. The average temperature during the experiments is 21° C.

To prepare for an experiment, we collect 1 kg of worker bees from the honey supers of five beehives in a 5 gallon bucket with a mesh lid. A caged (Glorybee plastic cage, #14343) queen bee is added to the bucket from a queen bank. To minimize strain on the bee colonies, we only collect worker bees from a particular hive and use each queen bee once every two weeks. We add one tablespoon of Pro Winter Patties to the bucket, and we spray the bees with sugar water once per day. This package of queen and workers is kept in the lab for three days to acclimate them to one another before beginning the experiment.

We build the swarm by attaching the queen bee to the center of the attachment surface, and then allowing the workers to swarm around her. After building the swarm, we leave it hanging in the laboratory for up for 3 - 4 days. We spray the swarm twice per day with sugar water, once two hours before any data collection, and once at the end of the day, and leave the bees several tablespoons of honey overnight. All room lights are turned off overnight. We run the x-ray for a maximum of three hours per day.

We find the weight of individual bees once for each swarm by freezing a sample of bees that is collected from the swarm after at least two hours have elapsed after feeding. Once the bees are frozen, we weigh a sample of 100 bees with an Escali scale (P115C) and find the average weight of a bee.

At the end of the swarm experiment, we return the queen bee to the queen bank. We coat the worker bees from the swarm in sugar powder and return them to the hives they were collected from. We wipe the swarm attachment board with ethanol before reusing it with a different swarm. We also sweep up any dead bees in the room, mop the floor, and wipe the x-ray emitter, filter, and detector with ethanol between swarm experiments.

#### 11.2 Calculation of bee grip strength

We measure the maximum force that a bee can support by tying a $153 \pm 11$ mm long elastic string (Bead Landing 0.5 mm elastic cord) to two bees, getting the bees to hold on to one another as in Supplementary Figure S1 **(a)**, and pulling them apart until they disconnect, as in Supplementary Figure S1 **(b)**. We record the trials with a smartphone camera and measure the initial length and final length of each elastic string and then use Hooke's law to measure the force between the bees. We find the spring constant by hanging masses with varying weights $w$ from 160 cm long pieces of the string and measuring the difference in final and initial length, and then plotting a linear best fit to the $L_f - L_0$ vs $w$ data from $0 \leq w \leq 71$ bee weights (Supplementary Figure S1 **(c)**). Since the initial length of the string is not constant in the bee trials (due to the bees moving around as we positioned the string), we calculate $k$ for each string length: $kL_{0calibration} = kL_{0bee}$. We perform a total of eight force measurement trials, each resulting in a top and bottom bee grip strength, excluding two in which the end of the string for the top bee was not visible in the camera image, for a total of 14 measurements of grip strength, $35 \pm 14$ bees.

#### 11.3 Experimental setup

In this section, we describe our procedure for setting up and taking CT scans. The experimental setup for the swarms during May - June 2021 had some small differences from the experimental setup in July - August 2021, which we describe here.

The experimental setup is shown in Supplementary Figure S2. We set up a swarm hanging from the attachment surface (A), a 33 cm diameter, 6 mm thick BCX wood disk. One side of the disk is smoother than the other, and we set up the swarm hanging from the rough side. This disk is attached to a hanging Everbilt 6-inch turntable bearing (B) using 4 - 6 6-32 nylon pan head screws with 6 mm diameter heads that protruded 2.5 mm from the attachment surface. Additionally, for the first two August trials, there are 3 and

4 TMP36 temperature sensors, respectively, with 4.6 mm diameter protruding 2.5 mm from the attachment surface. We plug any extra holes in the board with hot glue.

The other side of the turntable bearing is attached to a rectangular board with an attached stepper motor. The turntable is motorized by a Pololu Nema-17 stepper motor (Pololu #2267) controlled by a Pololu Tic T834 (May - June trials) or a Pololu Nema-23 stepper motor (Pololu #1477) (C) controlled with a Pololu Tic 36v4 (D) (July - August trials). The attachment surface and motorized turntable assembly hangs from an Ohaus Ranger 3000 scale (P31P15) (E) positioned on top a scissor jack (F) (July - August trials, for fine-tuning the height of the swarm).

To build the swarm, we remove the cage containing the queen from the bucket first and strap it to the center of the base board using 5 mm wide zip ties (J). We hang the experimental setup from the Ranger 3000 scale to measure the weight of the setup without the swarm. We then shake the bees into a bin underneath the caged queen. There is a stick sticking up from the bottom of the bin that barely touches the caged queen. The worker bees form a chain from the bin, up the stick, and to the swarm forming around the queen. As bees walk up the chain and join the swarm, we raise the queen and lower the bin to make room for additional bees on the swarm. Once the desired swarm weight is reached, we shake the remaining bees in the bin back into the bucket. We spray the swarm with sugar water and give it time to settle into its final shape for at least two hours before taking data.

The experimental setup is positioned in a JPI DynaVue: Digital X-Ray and Fluoroscopy in One System, between the x-ray emitter (G) and detector (H), $d = 17.7 \pm 0.7$ cm from the detector. The emitter and detector are fixed 90 cm apart. The detector has a 43 cm x 36 cm field of view, and the swarm is placed as close as possible to the detector to fit in the field of view. An 0.4 mm thick aluminum sheet is attached in front of the x-ray emitter as a filter to minimize beam hardening artifacts. We film the rotating swarm with a Logitech C270 HD webcam taped underneath the x-ray emitter (I).

A MATLAB script is used to control the stepper motor and continuously record its position and elapsed time every 0.1 s. The motor is set to continuously rotate and complete a full rotation in 25 seconds (16 pulses/step at 128 pulses/sec in May - June 2021, 256 pulses/step at 2048 pulses/sec in July - August 2021). This script additionally saves the weight of the swarm, calibration weight, distance from rotation axis to detector, x-ray settings, date and time, ambient temperature, and lab members present to a file.

The x-ray acquires projections in fluoroscopy mode at 15 frames per second, using 90-95 kV and 20 mA. Image acquisition is controlled with customized proprietary software by JPI. Projections are stored in raw .SDS files on a virtual drive, with approximately 370 projections corresponding to a full rotation of the swarm. A sample set of projections is shown in Supplementary video V1. To take a time-lapse of CT scans, we continuously revolve the swarm while saving projections. The size of a raw file is limited to 6 GB by the size of the virtual drive, so a continous recording of more than 3 revolutions requires more than one saved output file. The time stamps of all of the projections are stored in .html log files.

#### 11.4 CT scan processing

A script interpolates the motor position at each frame using the stored motor position over time and time stamps of saved frames and calculates the angle of each projection from the interpolated motor positions. We resize all of the projections to half the original size to speed up processing time. We then set the scale of each projection, $0.047 \pm 0.002$ cm/pixels, by measuring the diameter of the motor shaft. We obtain a background image by averaging 300 x-ray images taken without anything in front of the detector and then subtract this background from each projection to reduce noise.

The following paragrarphs describe the processing of the raw reconstructions, $\tilde{I}(x, y, z)$ to eliminate a cupping artifact. This is a known artifact in x-ray imaging, caused by beam hardening (18). During this processing, we also reduce noise that is caused by the short current and exposure time used to take the projections. Postproccesing to eliminate these artifacts results in a brightness matrix $I(x, y, z)$ that is linearly correlated to the mass of bees in each voxel.

We remove the noise around and inside the swarm in the reconstruction, as shown using a sample xy-slice from a 753-g swarm in Supplementary Figure S3. While we higlight the normalization and noise removal process with a single slice, this procedure is done to the entire 3D swarm reconstruction. The sample slice before processing is shown in Supplementary Figure S3(a).

1. Cropping and top-hat noise removal: We first measure the radius of the swarm board, and then detect the z-index of the start of the swarm near the board (characterized by a local minimum in the mean of the brightness of the reconstruction over the xy-plane). We then crop the reconstruction from that z-index to the swarm tip, $z = L$, and set the brightness outside of the swarm radius to zero. To reduce noise within the boundary of the swarm, we use a morphological top-hat filter on each xy-slice of the swarm (imtophat in MATLAB) with a 10-voxel radius disk structuring element. This performs a morphological opening of the image, which dilates and then erodes the image to remove small objects, and then subtracts the result from the original image, removing some of the uneven background noise within each slice. The sample slice after deleting the noise outside the swarm boundary and applying the top-hat filter is shown in Supplementary Figure S3(b). The cupping artifact is clearly visible in this slice - the center voxels are less bright than the voxels at the swarm boundary.

2. Boundary detection and brightness correction: Next, we correct the cupping artifact by normalizing the brightness at each voxel. We create a normalization coefficient matrix for the swarm. First we dilate the image (imdilate in MATLAB) with a 10-voxel radius sphere structuring element. We then sort the voxel values in this matrix from low to high brightness, and take the brightness value at the inflection point of the sorted values as a threshold. The dilated sample xy-slice is shown in Supplementary Figure S3(e). Voxels with a brightness below this threshold in the dilated image are outside of the swarm boundary. We save a separate logical matrix indicating whether a voxel is inside our outside the swarm boundary. We then remove edge effects at the boundary of the swarm. This is accomplished by dilating each slice with a 4-voxel radius disk filter and then setting the brightness in the entire region outside the swarm to the maximum brightness within that slice. The result from this is shown in Supplementary Figure S3(f). The final step in creating the normalization coefficient matrix is applying a 10-point Gaussian filter to blur sharp edges within the matrix. A sample xy-slice from the final normalization matrix is shown in Supplementary Figure S3(g). We then normalize the reconstruction values by dividing the reconstruction with top-hat filter applied, as in Supplementary Figure S3(b), by this normalization matrix. The swarm slice after normalization is shown in Supplementary Figure S3(c).

3. Constant-magnitude noise removal: Finally, we calculate the magnitude of the remaining noise within the swarm by separating the queen cage from the swarm and then performing a dilation on it with a structural element of a sphere with a 6-voxel radius. Since the cage contains only one bee and is otherwise occupied by empty space, we can take the minimum brightness within the dilated cage as the magnitude of the noise. Subtracting this noise from every voxel in the normalized swarm, as in Supplementary Figure S3(c), and then setting all the negative brightness values to zero leaves the queen cage with zero brightness everywhere except the queen bee herself. The final sample slice with noise removed is shown in Supplementary Figure S3(d).

After processing, we reject any reconstructions in which the outlines of the bees in the slices are not clearly defined. Upon paper acceptance, we will provide the CT data and analysis code in an external repository.

We measure the cupping artifact as the percentage by which the maximum brightness value within the bottom center 50 $\text{px}^3$ region differs from the maximum brightness of the entire swarm. The average cupping of the processed reconstructions is $7 \pm 5\%$, compared to $160 \pm 40\%$ for the raw reconstructions.

We set the scale of each reconstruction by measuring the side of the queen cage, 3.5 cm. The scale of the reconstructions is $s = 0.052 \pm 0.002$ cm/pixel.

#### 11.5 Prefactors of $W(z)$ and $M(z)$

The prefactor c in the expression for $\tilde{M}(z)$, with units of grams/cm, can be estimated as:

$$c = \overline{\rho} L^2 \tag{14}$$

where $L$ is the length of the swarm, and $\overline{\rho}$ is the mean swarm density calculated using the length of the side of each voxel, $s$, and the number of voxels in a layer that are within the swarm boundary, $N_B$:

$$\rho = \frac{1}{s^3 N_B} \sum_{x_i=-X/2}^{X/2} \sum_{y_i=-Y/2}^{Y/2} \sum_{z_i=0}^{L} m(x_i, y_i, z_i) \tag{15}$$

We can solve for the coefficient in the power law in Eq. 6 by factoring out $\tilde{M}(z)^a$ (as in Eq. 8 from the expression for $\tilde{W}(z)$ in Eq. 10 and comparing it to the power law in Eq. 6,

$$\tilde{C} = (\overline{\rho})^{1-a} g L^{3-2a} (1 - \frac{1}{a}). \tag{16}$$

We first calculate $\tilde{C}$ using the expression derived from the force balance, Eq. 16, which yields $\tilde{C} = 646 \pm 170$ $\mathrm{g}^{0.5}\mathrm{s}^{-2}\mathrm{cm}^{2.5}$. Second, we have a value for $C$ from the power law best fit to $W(z)$ vs. $M(z)$, Eq. 6, $C = 246 \pm 110$ $\mathrm{g}^{0.5}\mathrm{s}^{-2}\mathrm{cm}^{2.5}$. The units for $C$ are $\mathrm{g}^{0.5}\mathrm{s}^{-2}\mathrm{cm}^{2.5}$ only if $a = 1.5$. The deviation of $\tilde{C}$ from $C$ is $\frac{\tilde{C}-C}{\tilde{C}} = 0.63 \pm 0.08$.

The deviation $\tilde{C}$ from $C$, $\frac{\tilde{C}-C}{\tilde{C}} = 0.63 \pm 0.08$; we plot the deviation of $C$ from $\tilde{C}$ in Supplementary Figure S5**(c)**.

#### 11.6 Swarm dynamics

We show two sample slices from the swarm shown in **Figure 1(b-d)** in Supplementary Figure S4. In these slices, increased brightness correlates to higher mass at that voxel. Well-defined bright regions represent bees that were stationary over the 25 seconds of the scan, less bright regions are caused by moving bees not being fullly resolved due to motion blur, and dark regions represent empty space. The white rectangle in the top center of the xz-slice, Supplementary Figure S4(a), and in the center of the xy-slice, Supplementary Figure S4(b), is the queen cage, with the queen in the center of the cage in the xy-slice. We show a labeled close-up of a stationary bee in the inset of Supplementary Figure S4, consisting of a small circle, the thorax of a bee, and a longer ellipse, its abdomen. Stationary bees hang right side up, with their head and thorax above their abdomen. Bees are not evenly distributed throughout the swarm; rather, there appear to be regions near the top of the board and throughout the swarm where bees are more concentrated, with empty space around them.

We obtain snapshots of the arrangement of bees inside the swarm every 25 seconds by taking consecutive CT scans The xz-slices in Supplementary Video V3 shows that over the 11 minutes of this time lapse, the chains of bees remain in approximately the same positions, although individual bees may join or leave the chain. Other bees move around these stationary bees, resulting in a motion blur. This is demonstarted in the xy-slices in Supplementary Video V4, where the bees create a channel through which they can walk towards the queen. This is evidence that the channels hypothesized in (9) exist and serve an additional function to thermoregulation of providing pathways for bees throughout the swarm. How these channels change for thermoregulation and how the scaffolding might provide pathways through the swarm remains to be seen in future studies.
